# Mechanical communication through the ECM is frequency-dependent due to cell sensitivity to mechanical signal shape

**DOI:** 10.1101/2024.12.03.626471

**Authors:** Ido Nitsan, Stavit Drori, Shelly Tzlil

**Affiliations:** Faculty of Mechanical Engineering, Technion – Israel Institute of Technology, Haifa 32000 Israel

## Abstract

Cells communicate mechanically by sensing and responding to mechanical deformations generated by their neighbors in the extracellular matrix (ECM). By directly measuring the mechanical coupling between a cardiac cell and an artificial ‘mechanical cell’ and monitoring the dependence of beat-to-beat variability on mechanical coupling, we can quantify the sensitivity of cardiac cells to mechanical signals. Here we show that due to the dynamic viscoelastic properties of collagen hydrogels (a major component of the cardiac ECM), the shape of the mechanical signal changes in a frequency-dependent manner as it propagates through the gel, resulting in mechanical communication that depends on beating frequency. Moreover, we show that cardiac cell sensitivity to the shape of the mechanical signal results from its responsiveness to the loading rate, with an optimal loading rate for efficient mechanical communication. The dependence of signal shape on ECM viscoelasticity and the existence of an optimal loading rate suggest that there are ideal viscoelastic properties for mechanical communication between cardiac cells.

## Introduction

It was recently established that cells are able to communicate mechanically by responding to mechanical deformations generated by their neighbors (1–7). It is a challenging task to measure and define mechanical communication between cells. An additional challenge is to separate the mechanical component of intercellular communication from other modes of communication, such as a change in the amount or type of secreted chemo-attractants. Often, mechanical communication is defined as migration towards a contractile cell or frequency of cell-cell contact formation and mechanical coupling is only qualitatively estimated, for example, by imaging matrix fiber alignment spanning cell pairs (8, 9). In cardiac cells, we have shown previously that mechanical communication is essential for synchronized beating. We have previously shown that spontaneously beating cardiac cells can be paced using a ‘mechanical cell’, which consist of a tungsten probe that mimics the deformations generated by a neighboring beating cell (1). There is no physical contact between the cell and the probe, and the interaction therefore is purely mechanical and mediated through propagation of deformations in the matrix. In addition, we have measured mechanical coupling directly and showed that beat-to-beat variability decays exponentially with the level of mechanical coupling (3).

The ECM is highly non-linear viscoelastic material and therefore exhibits both elastic and viscous responses in a frequency dependent manner. In high frequencies it behaves as a pure elastic material while in lower frequencies molecular relaxation processes occur within the network and influence material response. Several recent studies have demonstrated the importance of ECM viscoelasticity in determining cell response (10–16). Intermediate viscosity was shown to maximize cell spreading (11) and fast stress relaxation results in robust cell migration on soft substrates (17).

Mechanical cell-cell communication is mediated by the extracellular matrix (ECM) and due to its viscoelasticity, cell-generated deformations will be filtered via the viscoelastic medium and therefore amplified, dissipated, or delayed in a frequency-dependent manner. As a result, the local mechanical signal generated by beating cardiac cells is expected to change its shape, phase, and amplitude in a way that depends on the beating frequency. Using collagen I, the major component of the cardiac ECM, we show that mechanical communication on collagen is frequency dependent. We demonstrate that this dependence results from changes in the mechanical signal shape as it propagates through the matrix. We further show that cell sensitivity to signal shape results from its sensitivity to the loading rate of the mechanical signal with an optimal loading rate of 1μm/sec at a beating frequency of 1Hz.

## Results and discussion

### Mechanical communication on ECM is frequency-dependent

Primary cardiomyocytes were isolated from neonatal rat hearts and cultured on either polyacrylamide gels or collagen I hydrogels. Hydrogels made of collagen I, a native component of the cardiac ECM, were used to explore the role of the dynamic viscoelastic properties of the ECM. We used glycated collagen matrices (18) with collagen concentration of 5mg/ml and ribose concentration of 250mM, to obtain Young’s modulus of ∼1 kPa, comparable to the Young’s modulus of our polyacrylamide substrate. Substrate stiffness in this range was shown to support optimal cardiac cell beating (19). We recently introduced the experimental setup (1, 3) which is illustrated schematically in Figure 1a. The ‘mechanical cell’ is a home-built mechanical device which consists of a tungsten probe mounted on a piezo stage with x-y-z actuators. Driven by the actuators, the probe cyclically pulls the substrate towards and away from the cardiac cell. The mechanical probe deforms the substrate such as to mimic the mechanical deformations generated by a neighboring beating cardiac cell. There is no physical contact between the cell and the probe and the interaction therefore is solely mechanical and mediated through propagation of deformations in the underlying substrate.

**Figure 1.**
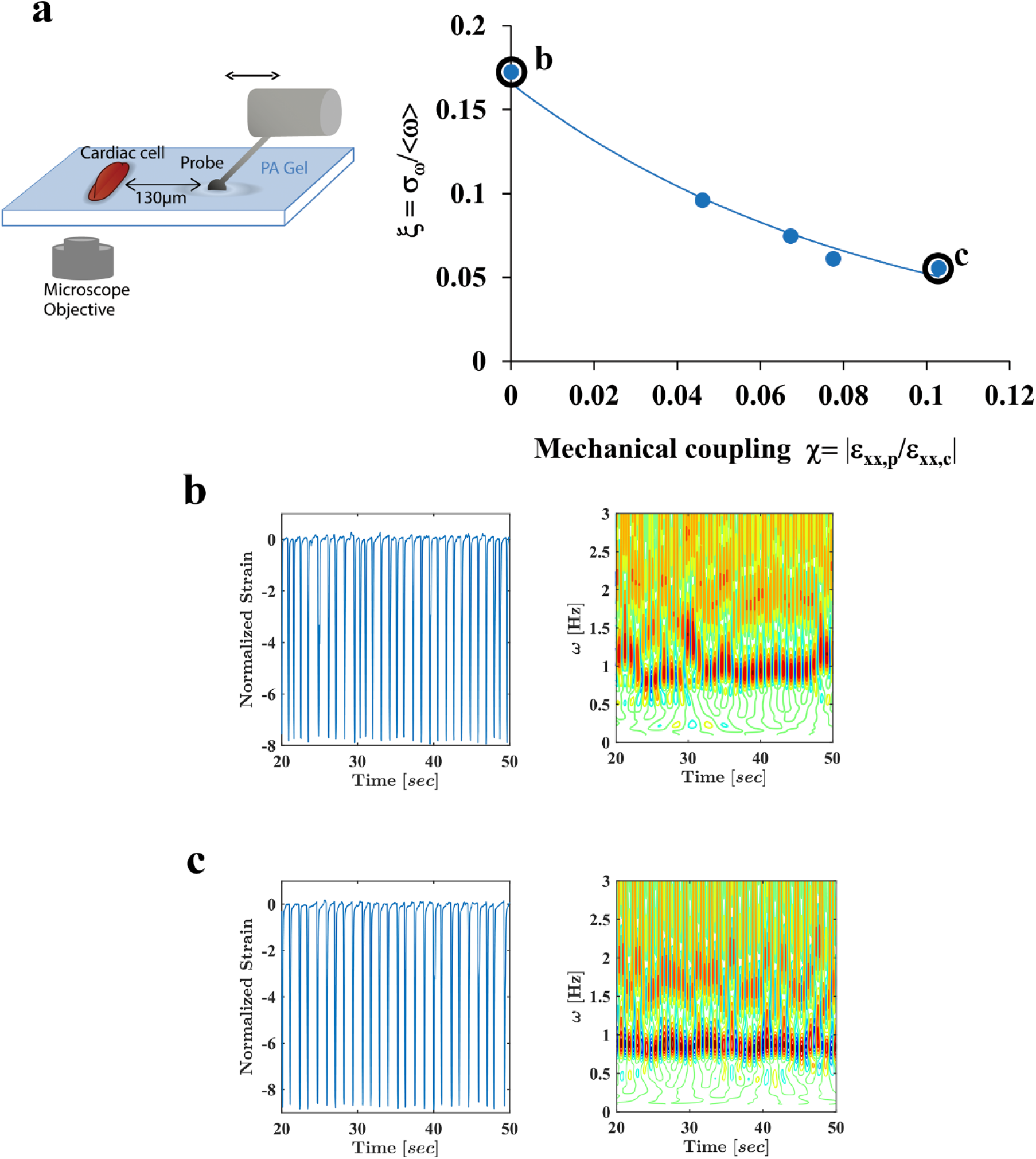
Beat-to-beat variability is reduced exponentially with increased mechanical coupling to a ‘mechanical cell’. a: *left:* A schematic representation of the ‘mechanical cell’ setup. A tungsten probe applies an oscillatory strain which mimics the deformations generated by a beating cardiac cell. *Right:* The dependence of beat-to-beat variability, for a representative isolated beating cardiac cell, on the strength of mechanical coupling to a ‘mechanical cell’. Beating noise (ξ) decays exponentially with the strength of mechanical coupling (*X*) to a ‘mechanical cell’ according to *ξ* = *ae*^−*kX*^ **b:** The normalized strain (*left*) and frequency (*right*) as a function of time without mechanical coupling, (the point marked in (a) by (b)). **c:** The normalized strain (*left*) and frequency (*right*) as a function of time for a mechanical coupling of *X*=0.103. (This point is marked in (a) by (c)).

We have recently shown that cardiac cells can be paced mechanically using a ‘mechanical cell’ (1) and that phase entrainment depends on the level of mechanical coupling (3). Isolated spontaneously beating cardiac cells have large beat-to-beat variability (variability in the time interval between consecutive contractions) that must be reduced to allow for synchronized contraction. In the intact heart, beat-to-beat variability serves as an indicator of cardiac health and increased beat-to-beat variability is associated with higher risks for cardiovascular events. We have recently shown that mechanical coupling regulates beat-to-beat variability (3).

By gradually increasing the amplitude of the ‘mechanical cell’ signal, we control the level of mechanical coupling. We directly measure the mechanical coupling of the cardiac cell to the ‘mechanical cell’ and monitor cardiac cell beat-to-beat variability as a function of mechanical coupling strength for the same cell. We let the cell interact with the probe for ten minutes before recording the signal and changing the amplitude to allow the cell to reach stationary behavior. Quantitatively, beat-to-beat variability of a cardiac cell (ξ) decays exponentially with the strength of mechanical coupling (*X*) to either a ‘mechanical cell’ or a neighboring cardiac cell according to ξ = *ae*^−*k*(ω)*X*^ where *k* is the exponential decay constant (Figure 1, Videos S1-S2 and (3)). The inverse value of the decay constant is the value of mechanical coupling required for the beat-to-beat variability to decay by a factor of 1/e. Therefore, *k*, can be used as a measure of cell sensitivity to mechanical coupling, or equivalently, as a measure of mechanical communication efficiency. Additional details on the experimental assay and the measurements of beat-to-beat variability and mechanical coupling can be found in the supplementary information (Fig. S1) and in previous work setup (1, 3).

In the general case, the exponential decay constant *k* is expected to depend on the beating frequency of the cardiac cell. However, as shown in Figure 2a-b, on linear elastic materials such as polyacrylamide, the exponential decay constant does not depend on frequency and equals 5.5±0.23. This indicates that on linear elastic materials, cell sensitivity to mechanical coupling is independent of beating frequency.

**Figure 2.**
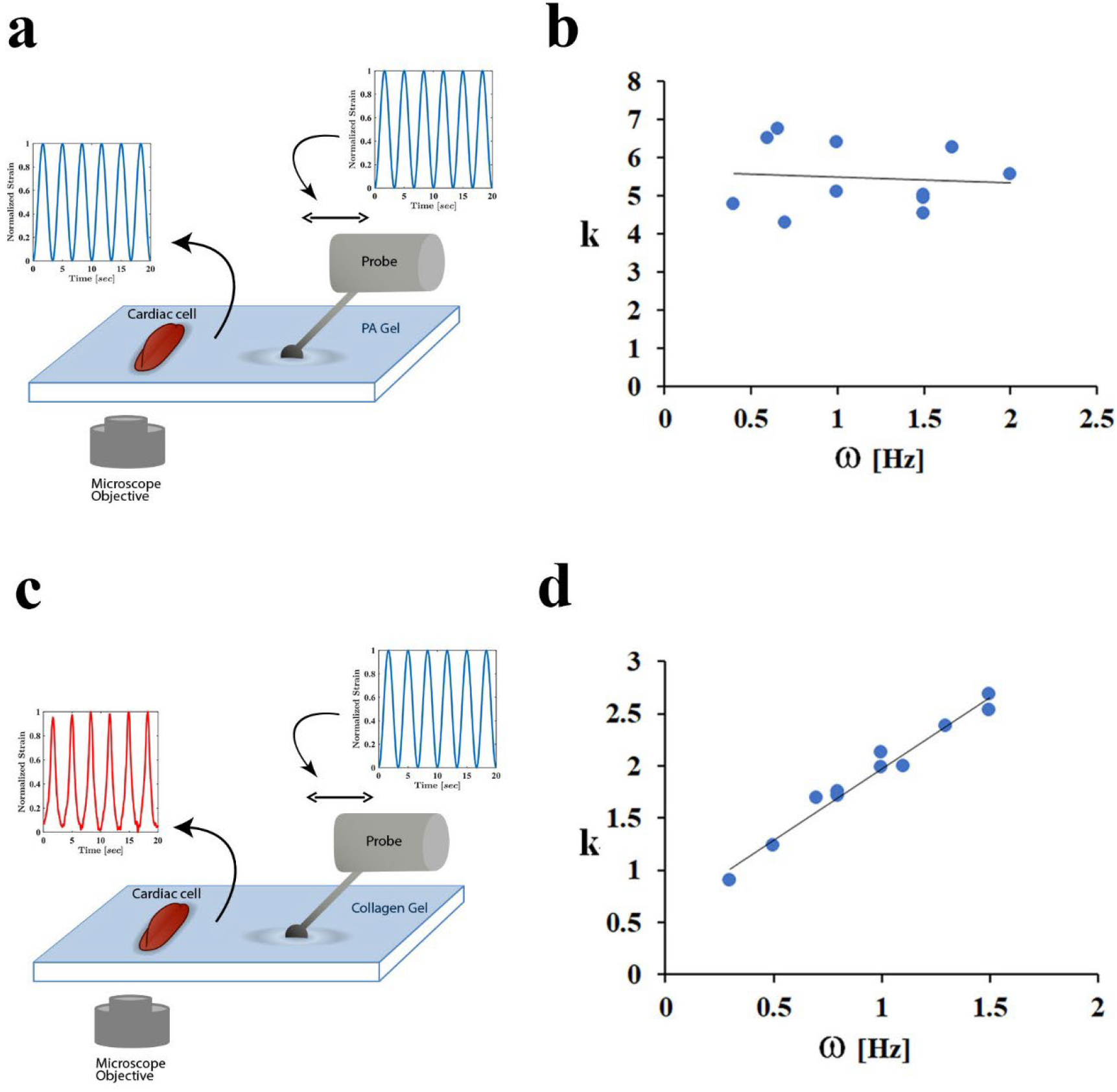
Mechanical communication is frequency-dependent on collagen ECM. **a,c:** Schematic illustration of the experimental protocols. An isolated cardiac cell interacts mechanically with a ‘mechanical cell’ on Polyacrylamide (PA) (a) or on collagen (c). The frequency of ‘mechanical cell’ oscillations is equal to the average frequency of the beating cardiac cell. A sinusoidal strain signal is applied by the mechanical cell and is measured next to the cell after it propagates through the substrate. On PA substrates (a) the signal still has sinusoidal form next to the cell (keeps its shape as it propagates through the gel). However, on collagen (c), the signal changes its shape as it propagates through the gel. **b**,**d:** Beat-to-beat variability (ξ) decays exponentially with the strength of mechanical coupling (*X*) to a ‘mechanical cell’ according to *ξ* = *ae*^−*k* (*ω* ) *X*^ (see Figure 1). The decay constant, *k*, is plotted as a function of frequency for isolated cardiac cells interacting with a ‘mechanical cell’ applying a sinusoidal mechanical signal on polyacrylamide (b) or collagen (d). Notice that the decay constant, *k* does not depend on frequency on polyacrylamide, while there is a strong dependence on frequency on collagen.

As shown in Figure 2c-d, for collagen hydrogels, the major component of the cardiac ECM, the behavior is qualitatively different. On collagen hydrogels, the decay constant, and therefore mechanical communication efficiency, is frequency-dependent and increases monotonically with the beating frequency within the examined range.

### Cells are sensitive to the shape of the mechanical signal

The nonlinearity and viscoelasticity of collagen are expected to change the shape of the oscillating mechanical signal as it propagates through the matrix. Indeed, a clear frequency-dependent change in the shape of the propagating strain was observed on collagen hydrogels (Figure 3a-b). Collagen fibers are unable to resist compressive strain and are known to buckle under cell-generated compressive forces and reorient along tensile forces (9, 20). Collagen buckling is observed in our experiments as shown in Fig. S2 and Video S2 and can explain the change in signal shape as it propagates through the collagen gel.

**Figure 3.**
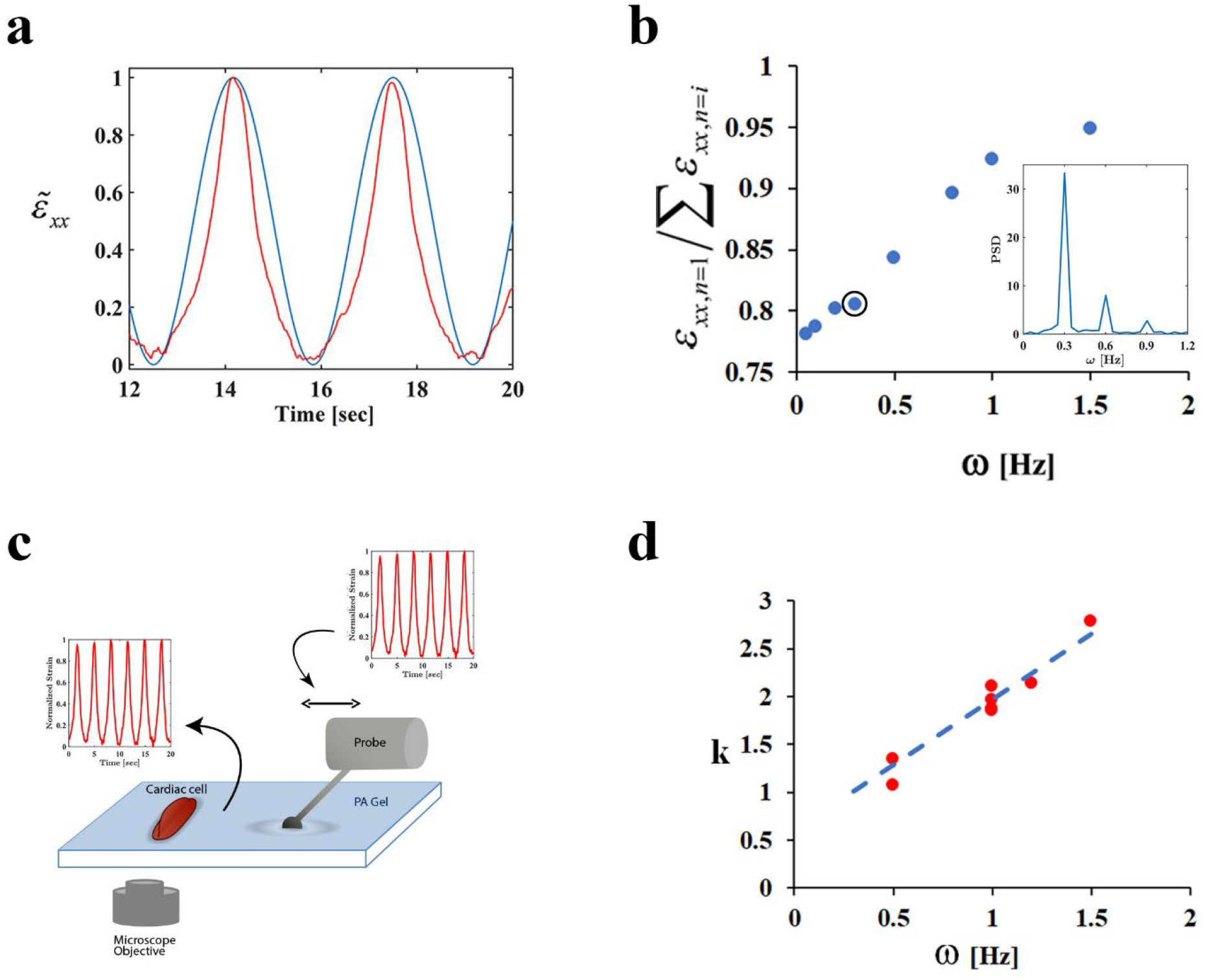
Cells are sensitive to the mechanical signal shape. **a:** Representative normalized strain signal at 0.3Hz next to the ‘mechanical cell’ (the location of force application, blue curve) and next to the cell, at a distance of d=84μm from the mechanical cell (after it propagates through the collagen gel,red curve). **b:** The weight of the first harmonic relative to the harmonic content as a function of frequency, at a distance of d=84μm from the mechanical cell. The marked point corresponds to the red signal in (a) and the corresponding Fourier transform is shown in the inset. **c**,**d:** Isolated cardiac cells interacting on polyacrylamide with an artificial mechanical cell that applies signals with the frequency-dependent shape observed on collagen next to the cell. c: Schematic illustration of the experimental protocol. d: Beat-to-beat variability (ξ) decays exponentially with the strength of mechanical coupling (*X*) to a ‘mechanical cell’ according to *ξ* = *ae*^−*k* (*ω* ) *X*^ (see Figure 1). The decay constant, *k*, as a function of frequency for the protocol schematically shown in (c) (red dots). The blue dashed line is the linear fit to the dependence of *k* on frequency for cells on collagen, presented in Figure 2d. Notice that the dependence of the decay constant, *k*, on frequency on <u>collagen</u> can be reproduced by mimicking on <u>polyacrylamide</u> the frequency-dependent signal shapes observed on collagen.

To test whether the change in shape of the mechanical signal is the origin of the frequency-dependent cell response, we repeated the experiment with a different protocol. As we have shown previously, since polyacrylamide gels are linear elastic gels, there is no apparent change in the shape of the signal while it propagates through the gel. Any signal shape (e.g., sinusoidal form) applied locally in the gel, will keep its form as it propagates in the gel and will have the same shape next to the cell (schematically shown in Figure 2a,3c). We repeat the experiment, but this time, for a given beating cell (e.g., with frequency ω_1_), we apply on the polyacrylamide gel an oscillatory signal that matches the shape of the signal observed on collagen at the same frequency (ω_1_) next to the cell (after it has propagated through the matrix). This experimental protocol is illustrated schematically in Figure 3c. For example, for a cell beating at 0.3Hz, the ‘mechanical cell’ generates on the polyacrylamide gel a signal with the shape of the red curve in Figure 2c rather than a sinusoidal signal.

As clearly shown in Figure 3d, we are able to reproduce the frequency-dependent cell response by mimicking on polyacrylamide the frequency-dependent signal shape observed on collagen. This implies that the frequency-dependent shape of the mechanical signal by itself can explain the change in cell sensitivity to mechanical coupling at different frequencies.

### Mechanical communication is most efficient at an optimal strain rate

As shown in Figure 3d, mechanical coupling efficiency depends on signal shape. Moreover, the signal shape by itself can reproduce the effect of having a viscoelastic material as a substrate. Different signal shapes are associated with different loading rates and it was recently shown that cell response is sensitive to the rate of applied force for fibroblast spreading under cyclic strain (21). To directly test the effect of loading rate on cell sensitivity to mechanical coupling and its ability to communicate mechanically efficiently, we repeat the experiments on polyacrylamide with a set of trapezoid signals. We used trapezoid signals since the loading rate is not constant for a sinusoidal signal whereas trapezoid signals have constant slopes and therefore a well-defined loading rate. All of the trapezoid strain profiles, have the same frequency but different slopes that correspond to different loading rates.

As done in the previous set of experiments, we gradually increase the trapezoid-shaped oscillations amplitude of the ‘mechanical cell’ at a frequency identical to the beating frequency of the cell and monitor beat-to-beat variability as a function of the strength of mechanical coupling. We conducted these experiments on cardiac cells that beat at 1Hz using 1Hz trapezoid signals on polyacrylamide substrates. When increasing the amplitude of the ‘mechanical cell’, we keep the loading rate of the trapezoid signal constant by changing the stall times between rise-and-fall.

As clearly shown in Figure 4a, the decay constant (*k*) of the beat-to-beat variability with mechanical coupling, i.e., the sensitivity of cells to mechanical coupling, has a non-monotonic dependence on the loading rate with a maximal value at a loading rate of ∼1μm/sec. Therefore, mechanical communication between cardiac cells beating at 1Hz is most efficient when the signal loading rate is ∼1μm/sec.

**Figure 4.**
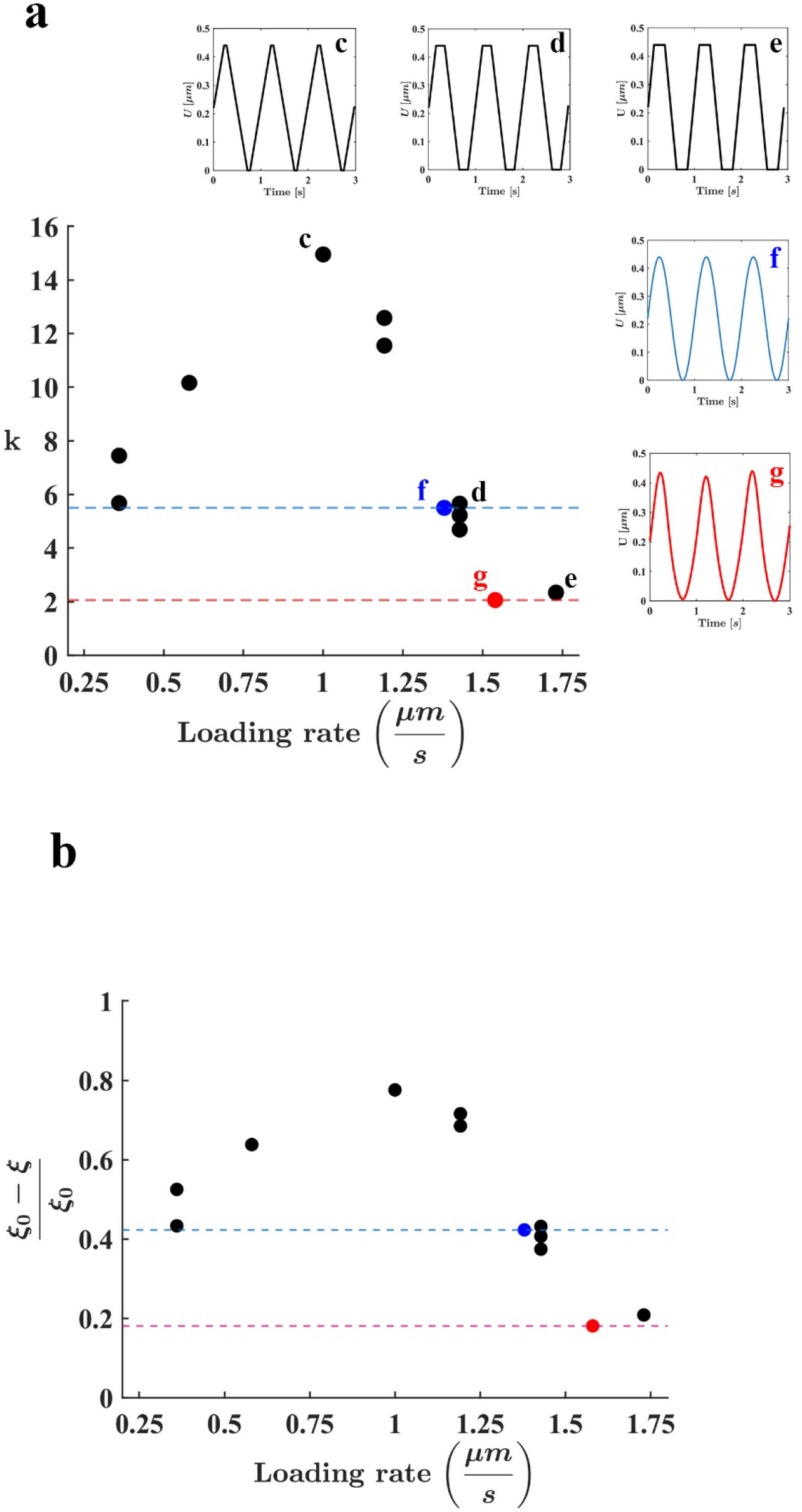
Mechanical communication is loading-rate dependent. **a:** Beat-to-beat variability (ξ) decays exponentially with the strength of mechanical coupling (*X*) to a ‘mechanical cell’ according to *ξ* = *ae*^−*k* (*ω* ) *X*^ (see Figure 1). The decay constant, *k*, which serves as a measure of cell sensitivity to mechanical coupling (or equivalently, mechanical communication efficiency) is plotted as a function of the loading rate of the mechanical signal for cells interacting with a trapezoid oscillatory signal (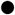, black circles), an oscillatory sinusoidal signal (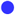, blue circle) and with an oscillatory signal shape observed on collagen next to the cell at 1Hz (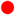, red circle). All signals are oscillatory signals applied on polyacrylamide at a frequency of 1Hz. For the non-trapezoid signals, the loading rate is defined as the maximal loading rate, i.e., the maximal derivative of the signal. **c-g:** the mechanical signals generated next to the cells in the points marked by **c-g** in the main graph respectively. k=5.5 and maximal loading rate=1.37 um/s for polyacrylamide; and k=2.1 and maximal loading rate=1.58 um/s for collagen. **b:** The change in beat-to-beat variability (ξ_0_-ξ) relative to the initial beat-to-beat variability (ξ_0_) as a function of the mechanical signal loading rate for mechanical coupling of *X*=0.1.

To incorporate the results obtained using oscillatory signals on polyacrylamide and collagen substrates at 1Hz (as reported in Figure 2b,d), we use the maximal loading rate of the mechanical signals, since the loading rate for these signals is not constant. As shown in Figure 4a, these points fall on the same curve.

As a reminder, *k* is the exponential decay constant of beat-to-beat variability as a function of mechanical coupling to a neighboring cell. To demonstrate how suboptimal loading rate, resulting in reduced mechanical coupling efficiency (i.e., a lower k) translates to increased beat-to-beat variability, we show the decrease in beat-to-beat variability as a function of loading rate for a given mechanical coupling of *χ*=0.1 (Figure 4b). This mechanical coupling strength (*χ*=0.1) corresponds, for example, to cells contracting at 5% strain on 3kPa PA separated by 80 μm along the axis perpendicular to the contraction direction, (equivalent to a distance of approximately 4-8 cell diameters). As clearly shown in Figure 4b, while beat-to-beat variability is reduced by nearly 80% in the optimal loading rate of 1 μm/sec, it is reduced by only 20% in the lowest or highest loading rates tested (1.73 μm/sec and 0.36 μm/sec respectively).

## Conclusions

Mechanical communication between cells is mediated by mechanical deformations which propagates through the ECM. Here we show that as the mechanical signal propagates through the ECM, its shape and consequently the associated loading rate is changing, thereby influencing mechanical communication. Modifications to the viscoelasticity of the mechanical environment (for example, by changing the level of enzymatic crosslinking in the ECM or matrix composition), will cause a change in the mechanical signal shape. The existence of an optimal loading rate for mechanical communication, therefore, suggests that there are optimal viscoelastic properties for mechanical communication.

Understanding the dependence of mechanical communication on the dynamic viscoelastic properties of the environment will allow us to tune ECM mechanics in order to achieve synchronized behavior of interacting cells or repair mechanical communication to restore normal function.

## Materials and Methods

All laboratory procedures conform to the Guide for the Care and Use of Laboratory Animals published by the U.S. National Institutes of Health. Animal usage was approved by the Animal Care and Use Committee of the Technion, Israel Institute of Technology.

### Cell culture

Neonatal cardiomyocytes were isolated from zero-day-old Sprague Dawley (SD) rat pups using the Neonatal Cardiomyocyte Isolation System (Worthington, NJ) according to the manufacturer’s instructions as described before(1). Briefly, hearts were rinsed with ice-cold HBSS and trypsinized for 16 hours at 4°C. Following trypsinization, hearts were incubated with trypsin inhibitor at 37°C, and then with collagenase solution for 30 minutes. Cells were triturated, pelleted and resuspended in L15 media (Worthington, NJ). Cells were then subjected to Percoll gradient for separation of cardiomyocytes from non-myocytes. Isolated cardiomyocytes were resuspended with culture media: F10 media (Sigma-Aaldrich, MO) supplemented with 5% fetal calf serum (Biological-Industries, Israel), 5% donor horse serum (Life-technologies, CA), penicillin 10u/ml, streptomycin 0.1mg/ml, 5-Bromo-2’-deoxyuridine (5-BrdU) 0.05mg/ml and CaCL_2_ 1mM (Signa-Aldrich, MO). Cultured cardiomyocytes were grown for at least 72 hours and up to 6 days.Approximately 8×10^5^ cells were plated on 30mm glass bottom plates (Greiner, Austria) covered with either collagen or matrigel patterned polyacrylamide gels.

### PDMS stamp: microfabrication and soft lithography

The patterns were designed using CleWin 5 (PhoeniX Software, NL). Isolated pairs of rectangles were designed. Rectangles were either 20 x 50um or 20 x 100um. Each rectangle represents a place that will be occupied by a cardiac cell. The distance between two cells in each pair varies throughout the mask (e.g. 10, 20, 40, 80, 120um). Chrome masks were manufactured by Delta Mask (Delta Masks BV, NL). A Si master with geometric patterns was fabricated using standard photolithographic technique as previously described(22). PDMS prepolymer was obtained via mixing Silicone elastomer and curing agent at 10:1 ratio (Dow Corning, Midland, MI) and degassing the mixture in a desiccator for 30 min. The prepolymer solution was poured over the Si master mold and cured in the oven at 80°C for 2 h. The elastomeric stamp was then peeled off carefully and cut in 1 × 1 cm^2^ squares for micropatterning.

### Intermediate patterned glass coverslips

Either Matrigel or laminin (Corning Inc, NY) was diluted 1:10 in L15 medium (Thermo Fisher). 160µl of the solution was added to the top of elastomeric microstamps, and incubated at 4°C overnight. Following incubation, the solution was aspirated, and the surface of the stamps was dried with a low stream of N_2_ gas. The stamps were then used to micropattern clean glass coverslips, via microcontact printing. The stamp was brought into complete conformal contact with a glass coverslip substrate for 5min at room temperature. Small weights (38 g) were placed over the PDMS stamp to aid complete protein pattern transfer from PDMS to intermediate glass

### Patterned polyacrylamide plates

Polyacrylamide (PA) gel coated plates were prepared following the protocol described previously(3). Briefly, 30mm glass bottom plates (Greiner, Austria) were activated using 2% 3-aminopropyltrimethoxysilane and 1% glutaraldehyde solution. Polyacrylamide/Bis-acrylamide 7.5/0.03% solution was mixed with 0.02% suspension of carboxylated dark red fluospheres (Life Technologies), 0.05% Ammonium Persulfate and 0.4% Temed (Bio-Rad, CA). 7.5 µl of mixed PA solution was applied on each activated plate and covered with, matrigel patterned glass coverslip, to form a thin film. The patterned matrigel on the coverslips was transferred to the surface of the polyacrylamide substrates during gelation. Films were left to polymerize for 30 min in room temperature. Following polymerization, polyacrylamide substrates were incubated in phosphate-buffered saline for at least 1 h and the top coverslips were carefully removed with a razor blade(23)

### Preparation of collagen coated dishes

Corning’s collagen I (HC) solution was glycated before gel formation. Collagen solution was mixed with ribose in 0.1% acetic acid to final concentration of 250mM ribose, and incubated for 5 days at 4°*C* as previously described(24). Glycated collagen gel coated plates were prepared according to manufacturer protocol (Corning’s coating procedure)(25). 30mm glass bottom plates (Greiner, Austria) were activated using 2% 3-aminopropyltrimethoxysilane, and 1% glutaraldehyde solution. 25 *μl* of glycated collagen solution was applied on each activated plate and covered with the 18mm round glass coverslip, to form a thin film. Films were incubated in 37°*C* 1 hour gelation.

### Labeling collagen fibers

Collagen fibers were labeled using NHS-Rhodamine in a mole ratio of 1:14, for 2 hours on ice. Non-reacted NHS-Rhodamine was removed by dialysis in 16.65M acetic acid

### Spinning disc confocal microscopy

Imaging was done using a Nikon Ti-E inverted microscope equipped with two Evolve EMCCD cameras (Photomatrix), 4 laser lines (405,491,561,642), phase contrast and Yokagawa spinning disc confocal. The dish was maintained at 37 ºC with 5% CO_2_/95% air using a cage incubator (Okolab). Imaging was done using 40x 0.95 NA (air) PlanApo Lambda objective (Nikon). Fluorospheres were excited using 642nm laser (Vortran) and Lifeact-RFP was excited with 561nm laser (Cobolt). Lifeact imaging was done using 100x 1.49 NA oil objective (Nikon).

### Mechanical probe assay

A rigid Tungsten probe with 25um tip diameter (Signatone) was mounted on a three axes (X-Y-Z) micro-positioning stages (Thorlabs) installed on a custom made adaptor for the microscope stage. The probe’s tip is immersed into the culture medium and slightly indents the PA substrate (2um indentation) 130um away from the target cell as previously described(1). By gradually increasing the amplitude of probe oscillations, the beating noise of a spontaneously beating cardiac cell can be monitored as a function of the strength of mechanical interaction with the mechanical probe. We let the cell interact with the probe for 10 minutes before recording the signal and changing the amplitude to allow the cell to reach stationary behavior. Probe oscillation frequency is chosen to be equal to cell frequency in order to separate the effect of pacing from that of noise reduction.

## Supplementary information

is available.

## Acknowledgements

This work was supported by the Israel Science Foundation grants 507/18 and 1009/23.

## Author contributions

S.T designed research and wrote the manuscript. I.N. and S.D. performed experiments, I.N and S.T. analyzed the data, S.T. supervised the research. All authors discussed and helped prepare the manuscript.

## Competing interests

The authors declare no competing interests

## Notes

### Competing Interest Statement

The authors have declared no competing interest.

### Summary of Updates

The figures were rearranged to clarify the results. The buckling behavior of collagen was moved to the supplementary information.

## References

1. I. Nitsan, S. Drori, Y. E. Lewis, S. Cohen, S. Tzlil, Mechanical communication in cardiac cell synchronized beating. Nat Phys 12, 472–477 (2016). DOI:10.1038/nphys3619.

2. T. E. Angelini, E. Hannezo, X. Trepat, J. J. Fredberg, D. A. Weitz, Cell Migration Driven by Cooperative Substrate Deformation Patterns. Phys Rev Lett 104, 168104 (2010).

3. H. Viner, I. Nitsan, L. Sapir, S. Drori, S. Tzlil, Mechanical Communication Acts as a Noise Filter. iScience 14 (2019). DOI:10.1016/j.isci.2019.02.030.

4. C. A. Reinhart-King, M. Dembo, D. A. Hammer, Cell-Cell Mechanical Communication through Compliant Substrates. Biophys J 95, 6044–6051 (2008). DOI:10.1529/biophysj.107.127662.

5. K. K. Chiou, J. W. Rocks, C. Y. Chen, S. Cho, K. E. Merkus, A. Rajaratnam, P. Robison, M. Tewari, K. Vogel, S. F. Majkut, B. L. Prosser, D. E. Discher, A. J. Liu, Mechanical signaling coordinates the embryonic heartbeat. PNAS 113, 8939–8944 (2016). DOI:10.1073/pnas.1520428113.

6. C. Yang, X. Dong, B. Sun, T. Cao, R. Xie, Y. Zhang, Z. Yang, J. Huang, Y. Lu, M. Li, X. Wang, Y. Xu, F. Ye, Q. Fan, Physical immune escape: Weakened mechanical communication leads to escape of metastatic colorectal carcinoma cells from macrophages. Proceedings of the National Academy of Sciences 121, e2322479121 (2024). DOI:10.1073/PNAS.2322479121.

7. X. Tang, P. Bajaj, R. Bashir, T. A. Saif, How far cardiac cells can see each other mechanically. Soft Matter 7, 6151–6158 (2011). DOI:10.1039/c0sm01453b.

8. C. D. Davidson, F. S. Midekssa, S. J. DePalma, J. L. Kamen, W. Y. Wang, D. K. P. Jayco, M. E. Wieger, B. M. Baker, Mechanical Intercellular Communication via Matrix-Borne Cell Force Transmission During Vascular Network Formation. Advanced Science 11 (2024). DOI:10.1002/advs.202306210.

9. X. Chen, D. Chen, E. Ban, K. C. Toussaint, P. A. Janmey, R. G. Wells, V. B. Shenoy, Glycosaminoglycans modulate long-range mechanical communication between cells in collagen networks. Proceedings of the National Academy of Sciences of the United States of America 119 (2022). DOI:10.1073/PNAS.2116718119/FORMAT/EPUB.

10. O. Chaudhuri, L. Gu, D. Klumpers, M. Darnell, S. A. Bencherif, J. C. Weaver, N. Huebsch, H.-P. Lee, E. Lippens, G. N. Duda, D. J. Mooney, Hydrogels with tunable stress relaxation regulate stem cell fate and activity. Nature Materials 15, 326–334 (2016). DOI:10.1038/NMAT4489.

11. Z. Gong, S. E. Szczesny, S. R. Caliari, E. E. Charrier, O. Chaudhuri, X. Cao, Y. Lin, R. L. Mauck, P. Janmey, J. A. Burdick, V. B. Shenoy, Matching material and cellular timescales maximizes cell spreading on viscoelastic substrates. Proceedings of the National Academy of Sciences of the United States of America 115, E2686–E2695 (2018). DOI:10.1073/PNAS.1716620115/SUPPL_FILE/PNAS.201716620SI.PDF.

12. O. Chaudhuri, L. Gu, M. Darnell, D. Klumpers, S. A. Bencherif, J. C. Weaver, N. Huebsch, D. J. Mooney, Substrate stress relaxation regulates cell spreading. Nature Communications 6 (2015). DOI:10.1038/ncomms7365.

13. O. Chaudhuri, J. Cooper-White, P. A. Janmey, D. J. Mooney, V. B. Shenoy, Effects of extracellular matrix viscoelasticity on cellular behaviour. Nature 2020 584:7822 584, 535–546 (2020). DOI:10.1038/s41586-020-2612-2.

14. A. Elosegui-Artola, A. Gupta, A. J. Najibi, B. R. Seo, R. Garry, C. M. Tringides, I. de Lázaro, M. Darnell, W. Gu, Q. Zhou, D. A. Weitz, L. Mahadevan, D. J. Mooney, Matrix viscoelasticity controls spatiotemporal tissue organization. Nature Materials 22 (2023). DOI:10.1038/s41563-022-01400-4.

15. W. Fan, et al., Matrix viscoelasticity promotes liver cancer progression in the pre-cirrhotic liver. Nature 626 (2024). DOI:10.1038/s41586-023-06991-9.

16. J. P. Winer, S. Oake, P. A. Janmey, Non-Linear Elasticity of Extracellular Matrices Enables Contractile Cells to Communicate Local Position and Orientation. PLoS ONE 4, e6382 (2009). DOI:10.1371/journal.pone.0006382.

17. K. Adebowale, Z. Gong, J. C. Hou, K. M. Wisdom, D. Garbett, H. Lee, S. Nam, T. Meyer, D. J. Odde, V. B. Shenoy, O. Chaudhuri, Enhanced substrate stress relaxation promotes filopodia-mediated cell migration. Nature Materials 2021 20:9 20, 1290–1299 (2021). DOI:10.1038/s41563-021-00981-w.

18. M. S. Hall, F. Alisafaei, E. Ban, X. Feng, C.-Y. Hui, V. B. Shenoy, M. Wu, Fibrous nonlinear elasticity enables positive mechanical feedback between cells and ECMs. Proceedings of the National Academy of Sciences of the United States of America 113, 14043–14048 (2016). DOI:10.1073/pnas.1613058113.

19. A. J. Engler, C. Carag-Krieger, C. P. Johnson, M. Raab, H. Y. Tang, D. W. Speicher, J. W. Sanger, J. M. Sanger, D. E. Discher, Embryonic cardiomyocytes beat best on a matrix with heart-like elasticity: scar-like rigidity inhibits beating. J Cell Sci 121, 3794–3802 (2008).

20. Y. L. Han, P. Ronceray, G. Xu, A. Malandrino, R. D. Kamm, M. Lenz, C. P. Broedersz, M. Guo, Cell contraction induces long-ranged stress stiffening in the extracellular matrix. Proceedings of the National Academy of Sciences 115, 4075–4080 (2018). DOI:10.1073/PNAS.1722619115.

21. I. Andreu, B. Falcones, S. Hurst, N. Chahare, X. Quiroga, A.-L. Le Roux, Z. Kechagia, A. E. M. Beedle, A. Elosegui-Artola, X. Trepat, R. Farré, T. Betz, I. Almendros, P. Roca-Cusachs, The force loading rate drives cell mechanosensing through both reinforcement and cytoskeletal softening. Nature Communications 2021 12:1 12, 1–12 (2021). DOI:10.1038/s41467-021-24383-3.

22. X. Tang, M. Yakut Ali, M. T. A. Saif, A novel technique for micro-patterning proteins and cells on polyacrylamide gels. Soft Matter 8, 7197 (2012). DOI:10.1039/c2sm25533b.

23. A. J. S. Ribeiro, Y.-S. Ang, J.-D. Fu, R. N. Rivas, T. M. A. Mohamed, G. C. Higgs, D. Srivastava, B. L. Pruitt, Contractility of single cardiomyocytes differentiated from pluripotent stem cells depends on physiological shape and substrate stiffness. Proceedings of the National Academy of Sciences 112, 12705–12710 (2015). DOI:10.1073/pnas.1508073112.

24. R. Roy, A. Boskey, L. J. Bonassar, Processing of type I collagen gels using nonenzymatic glycation. J Biomed Mater Res A 9999A, NA-NA (2009). DOI:10.1002/jbm.a.32231.

25. Corning collagen I coating procedures, <https://certs-ecatalog.corning.com/life-sciences/certs/354249/354249_9343002.pdf> (October 11, 2022).

