## Supplementary information for "Mechanical communication through the ECM is frequency-dependent due to cell sensitivity to mechanical signal shape"

**1. Measuring beat-to-beat variability and mechanical coupling:**

The beating noise (beat-to-beat variability) $\xi$ is defined as the relative standard deviation of the beating frequency, $\xi_\omega = \sigma_\omega / < \omega >$ , where $< \omega >$ is the average beating frequency of the cell and $\sigma_\omega$ is the standard deviation.

Cells sense the deformation (strain field) at their edge, which are the sum of their own generated deformations and those generated by nearby beating cells. A neighboring cell is expected to have a significant influence when its deformations are comparable to or higher than those generated by the cell itself. Therefore, as explained in detail in (1) and demonstrated in Figure S1 below, mechanical coupling is defined as the absolute value of the ratio between the strain generated by a neighboring cell/'mechanical cell' at the edge of the cell over the strain generated by the cell itself. More formally, we define a mechanical coupling parameter $\chi$, such that $\chi = \left| \varepsilon_{xx,n} / \varepsilon_{xx,c} \right|$ where $\varepsilon_{xx,n}$ is the strain generated by a neighboring cell/'mechanical cell' along the vector connecting the two cells (x-axis) and $\varepsilon_{xx,c}$, is the strain generated by the cell itself. This ratio defines how strong is the perturbation generated by the neighboring cell/ 'mechanical cell'.

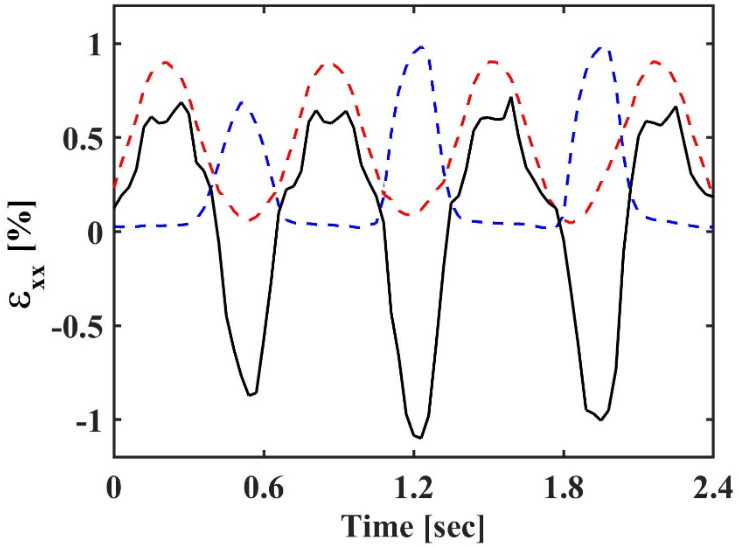


**Figure S1: Measurement of mechanical coupling between a beating cardiac cell and a 'mechanical cell':** The average strain at the edge of the beating cell in the direction perpendicular to the contraction axis (black curve) is generated by both the cardiac cell and the mechanical cell. The blue and red dashed curves are the normalized signals of the beating cell and the 'mechanical cell' respectively, during a short time interval where the cell and the 'mechanical cell' beat in anti-phase. The normalized signals are shown in order to determine the relative contributions of the beating cell and the 'mechanical cell' to the total strain. The mechanical coupling is defined as the absolute value of the ratio between the strain generated by the 'mechanical cell' at the edge of the cell over the strain generated by the cell itself at the same location, $\chi = \left| \varepsilon_{xx,n} / \varepsilon_{xx,c} \right|$.

### 2. Collagen buckling in mechanical communication

Collagen fibers are unable to resist compressive strain and are known to buckle under compressive forces generated by cells. Collagen buckling is observed in our experiments (Fig. S2 below and Movie S2) and can explain the changes in mechanical signal shape as it propagates through collagen gels.

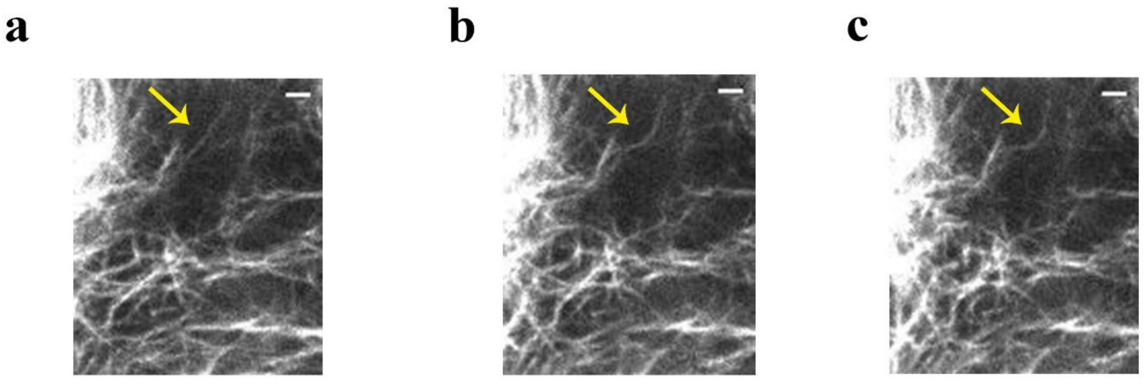


**Figure S2: Collagen buckling in mechanical communication:. a-c:** Representative movie frames (MovieS2) of labeled collagen fibers locally deformed by the 'mechanical cell' at three time points: beginning of 'mechanical cell' movement (**a**, time zero, no deformation), after 1.5 seconds (**b**) and after 1.8 seconds (**c**). The yellow arrow is pointing at a representative fiber which undergoes buckling. Scale bar is: 5μm.

### 3. Movie Legends

**MovieS1: Bead displacement generated by an isolated beating cardiomyocyte:** Time lapse imaging of the displacement field generated by an isolated beating cardiac cell on a polyacrylamide substrate. Mechanical deformations are detected by following the displacement of 0.2mm fluorescent beads embedded within the gel. The corresponding beat-to-beat variability is shown in Figure 1b in the main text. The movie is played in real time. Scale bar is 10μm

**MovieS2: Bead displacement generated by a beating cardiomyocyte interacting with a 'mechanical cell':** Time lapse imaging of the displacement field generated by the same cardiac cell shown in MovieS1 on a polyacrylamide substrate, now interacting with a mechanical cell. The corresponding beat-to-beat variability is shown in Figure 1c in the main text. The movie is played in real time. Scale bar is 10μm.

**MovieS3: Collagen fibers buckling under 'mechanical cell' oscillations:** Time lapse imaging of collagen fibers under oscillatory forces applied by a 'mechanical cell'. Collagen fibers are labeled using NHS-Rhodamine. Snapshots from the movie are shown in Figure S2. The movie is played in real time. Scale bar is 5μm.

### References:

1. H. Viner, I. Nitsan, L. Sapir, S. Drori, S. Tzlil, Mechanical Communication Acts as a Noise Filter. *iScience* **14** (2019).
